# A first assessment of the distribution and abundance of large pelagic species at Cocos Ridge seamounts (Eastern Tropical Pacific) using drifting pelagic baited remote cameras

**DOI:** 10.1101/2020.12.09.417493

**Authors:** Marta Cambra, Frida Lara-Lizardi, Cesar Peñaherrera, Alex Hearn, James T. Ketchum, Patricia Zarate, Carlos Chacón, Jenifer Suárez-Moncada, Esteban Herrera, Mario Espinoza

## Abstract

Understanding the link between seamounts and large pelagic species (LPS) is critical for guiding management and conservation efforts in open water ecosystems. The seamounts along the Cocos Ridge in the Eastern Tropical Pacific (ETP) are thought to play a critical role for LPS moving between Cocos Island (Costa Rica) and Galapagos Islands (Ecuador). However, to date, research efforts to understand pelagic community structure beyond the borders of these oceanic Marine Protected Areas (MPAs) have been limited. This study used drifting-pelagic baited remote underwater video stations (BRUVS) to characterize the distribution and relative abundance of LPS at Cocos Ridge seamounts. Our drifting-pelagic BRUVS detected a total of 21 species including sharks, large teleosts, small teleosts, dolphins and one sea turtle, of which 4 are threatened species. Relative abundance and richness of LPS was significantly higher at shallow seamounts (<400m) compared to deeper ones (>400m) suggesting that seamount depth could be an important driver structuring LPS assemblages along the Cocos Ridge. Our cameras provided the first visual evidence of the schooling behaviour of *S. lewini* at two shallow seamounts outside the protection limits of Cocos and Galapagos Islands. However, further research is still needed to demonstrate a positive association between LPS and Cocos Ridge seamounts. Our findings showed that drifting pelagic BRUVS are an effective tool to survey LPS in fully pelagic ecosystems of the ETP. This study represents the first step towards the standardization of this technique throughout the region.

## Introduction

Quantifying the spatial distribution and abundance of pelagic species is critical to effectively manage and protect their populations in open oceans [1]. Overexploitation of these highly productive regions is driving many large pelagic species (LPS) such as elasmobranchs, large teleosts, sea turtles and cetaceans to dangerously low levels [2,3], raising global concerns about the potential top-downs effects on marine ecosystems [4,5]. Seamounts have been recognized as productive and unique features in open water-systems where highly migratory LPS tend to aggregate, thus becoming highly vulnerable areas to overfishing [6,7]. Understanding the link between seamounts and LPS is critical for identifying regional hotspots of biological production, and guiding management and conservation efforts in open water ecosystems [8].

The Cocos Ridge is a chain of seamounts in the Eastern Tropical Pacific (ETP) that connects Cocos Island (Costa Rica) and the Galapagos archipelago (Ecuador) [9]. These two oceanic island groups are considered biodiversity hotspots in the ETP because of the high biomass of apex predators they harbor [10,11]. They are also Marine Protected Areas (MPAs) and UNESCO World Heritage Sites [12,13]. Previous studies have shown a higher degree of movement connectivity between Cocos and Galapagos Islands relative to other regions of the ETP, suggesting that LPS may be using this area as a migratory corridor [14–18]. Cocos Ridge seamounts are thought to be sites of ecological importance where LPS aggregate during these migrations [18]. However, most research effort on LPS in the ETP have concentrated inside MPAs [19–22] and therefore, the biological diversity and community structure associated to seamounts have still not been described.

As LPS continue decreasing in the ETP [19,21,23], there is a greater need to survey the open ocean to effectively guide marine spatial planning [24]. Information outside the protection boundaries of Cocos and Galapagos Islands is scarce and restricted to fishery dependent data [25–27] or to movement studies on sharks [14–17,28], teleosts [29] and sea turtles [30,31]. Despite the valuable information acquired from biotelemetry devices to understand individual habitat preferences, movements and migrations, this technique relies on the catch of a high number of individuals from various species in order to understand pelagic community structure and identify aggregation sites in the open ocean [32]. Furthermore, such studies can be invasive, expensive and logistically challenging [33]. Fisheries data also present some limitations as they are usually biased by temporally and spatially uneven sampling effort, gear selectivity and lack of robust reports [34,35].

Drifting-pelagic baited remote underwater video stations (BRUVS) have demonstrated a promising potential for studying pelagic wildlife in open water ecosystems [36–38]. Drifting-pelagic BRUVS are an adaptation of the benthic BRUVS where an anchoring system is no longer needed, thus enabling dynamic sampling over deep and topographically complex pelagic areas [37]. The odor of the bait triggers bait-search behavior in nearby fish assemblages, increasing the probability of detecting predatory species in the vicinity of the BRUVS [39]. A reduced amount of zeros derived from bait use increases the statistical power of BRUVS compared to traditional survey techniques [40–42]. Although the drifting-pelagic version of BRUVS has received little attention to date [43], it offers a powerful framework to overcome the difficulties associated to effectively survey pelagic assemblages [36]. For example, drifting-pelagic BRUVS units can be simultaneously deployed reducing the survey effort while generating permanent high definition images on species composition, behavior and relative abundance at different depth levels and for long periods of time [37]. Additionally, they are affordable and easy to operate allowing the participation of stakeholders into the field work. In this study, we used drifting-pelagic BRUVS for the first time in the ETP to characterize the distribution and relative abundance of LPS at Cocos Ridge seamounts. This study might serve as an important reference on the future use of drifting-pelagic BRUVS at a regional level.

## Methodology

### Study area

The Cocos Ridge is an underwater mountain range located in the northwestern Panama basin of the ETP, which originated more than 30 million years ago as a result of volcanic activity from the Galapagos Ridge hot spot [9]. The Cocos Ridge rises about 2000 m above the seafloor and extends more than 1000 km from the Galapagos Islands to Cocos Island, and from Cocos Island to the south pacific coast of Costa Rica [9,44,45]. Although the depth and total number of seamounts along the Cocos Ridge is not available on global bathymetric databases, there are at least 14 seamounts that have been identified in this region [18,44]. All the seamounts along the Cocos Ridge are located within the 200 Nm mile Exclusive Economic Zones (EEZs) of either Ecuador or Costa Rica (Fig 1).

**Fig 1.**
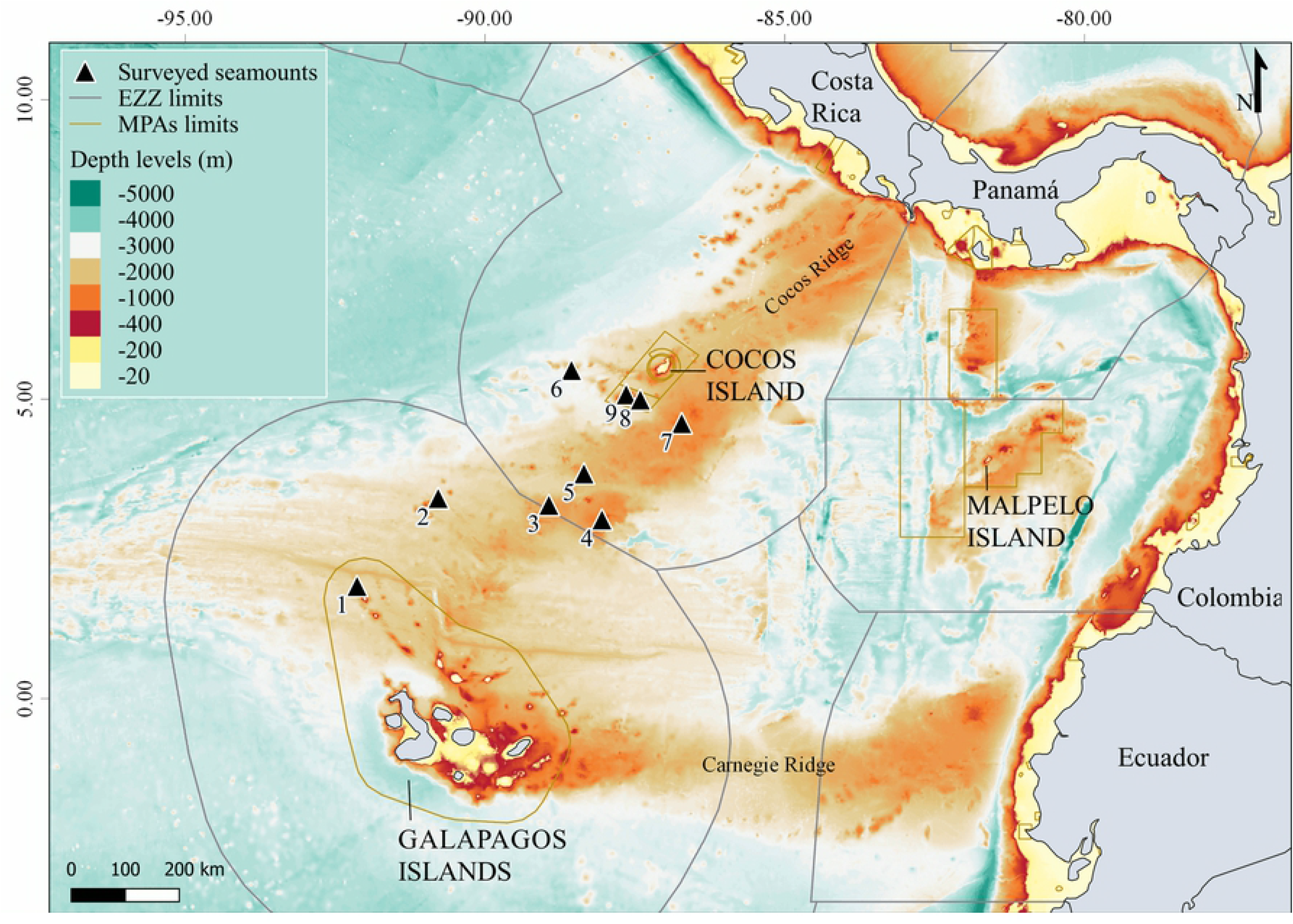
Location of seamounts surveyed along the Cocos Ridge between Cocos and the Galapagos Islands. Numbers indicate surveyed seamounts: 1) NW Darwin; 2) Paramount; 3) Medina 1; 4) Medina 2; 5) Medina 3; 6) West Cocos; 7) East Cocos; 8) Las Gemelas 1; 9) Las Gemelas 2. Bathymetry data was obtained from ETOPO1 1 Arc-Minute Global Relief Model [46].

The variable and dynamic oceanographic conditions surrounding the waters of Cocos Ridge seamounts are mainly attributed to the southern oscillation of the Intertropical Convergence Zone, which is also associated to seasonal changes on wind patterns and the convergence of different ocean currents [18,47]. There are several major oceanic current systems converging in this region, resulting in a unique variety of tropical and temperate marine life [48]. The North Equatorial Current (east – west), the South Equatorial Current (east – west) and the Equatorial Countercurrent (west – east) are the main surface currents affecting Cocos Ridge seamounts [49]. Although the central region of the ETP is characterized by warm waters (average 27.5°C), the Humboldt Current (the southern limit of the ETP) can bring cooler waters (~18°C) when the Intertropical Convergence Zone moves to the north, potentially affecting seamounts closer to Galapagos Islands. Oceanographic variability is accentuated by the increase of the average sea temperature during the El Niño–Southern Oscillation Phenomenon, which occurs at irregular intervals of 2 – 7 years [50]. Chlorophyll-A concentrations along the marine corridor between Cocos and Galapagos Islands respond to seasonal variations typically oscillating between 0.15 and 0.22 mg m^−3^ [18]. The ETP is also characterized by a shallow thermocline (often at 25 m) above a permanent barrier of cold hypoxic water that limits the available physical habitat for predator species [51].

### Sampling method

Nine seamounts along the Cocos Ridge were surveyed in April 2018 (Fig 1). Seamount selection was based on the depth of each seamount summit and available time in the field. We prioritized the shallowest seamounts we could detect in the study area to increase the probability of LPS detection rates in subsurface waters (10 and 25 m deep) (Table 1). The depth of each seamount was previously researched using the Seamount Catalog of EarthRef [52] and the National Center for Environmental Information of the National Oceanic and Atmospheric Administration [53]. The depth sounder of the research vessel used during the expedition provided more accurate locations for seamounts shallower than 600 m (depth sounder maximum capacity) (Table 1). Additional information regarding seamount bathymetry was accessed during the expedition from boat navigation charts and from offline nautical charts [54].

**Table 1.** Survey effort, location, depth and environmental data associated with seamounts surveyed along the Cocos Ridge.

| Seamounts | N | Hours | Latitude | Longitude | Depth | Temp | Drift | Chl-a |
| --- | --- | --- | --- | --- | --- | --- | --- | --- |
| Las Gemelas 2 | 9 | 17.5 | 5.06889 | -87.63444 | -172* | 28.1 / 26.8 | 0.65 | 0.18 |
| Las Gemelas 1 | 20 | 39.9 | 4.98500 | -87.40167 | -198* | 29 / 27.9 | 0.86 | 0.21 |
| East Cocos | 20 | 31.3 | 4.60000 | -86.72000 | -775 | 29.7 / 28.5 | 0.97 | 0.16 |
| West Cocos | 20 | 55.1 | 5.46750 | -88.53333 | -283* | 29.3 / 29.1 | 0.47 | 0.13 |
| Medina 3 | 18 | 26.5 | 3.33389 | -88.26806 | -445* | 28.7 / 27.2 | 2.19 | 0.21 |
| Medina 2 | 15 | 38.4 | 2.99000 | -88.05000 | -688 | 28.2 / 25.8 | 3.08 | 0.28 |
| Medina 1 | 19 | 29.8 | 3.23500 | -88.93500 | -818 | 28.9 / 27.2 | 1.91 | 0.31 |
| Paramount | 19 | 43.1 | 3.34900 | -90.78100 | -188* | 29.5 / 25.9 | 1.11 | 0.14 |
| NW Darwin | 10 | 11.7 | 1.88123 | -92.13404 | -1200 | 26.5 / 22.8 | 1.95 | 0.20 |

N – number of deployments; Hours –recorded hours with baited remote underwater video
stations (BRUVS); Depth (m) of seamount summit (\*): depth recorded in the field; Temp – mean temperature (°C) recorded on shallow (10 m) and deep (25 m) deployments; Drift – mean drifting speed (km h<sup>-1</sup>) of pelagic-BRUVS deployments; Chl-a – mean concentration of chlorophyll-a (mg m<sup>-3</sup>). Seamounts are ordered with increasing distance from Cocos Island.

The drifting-pelagic BRUVS used in this study are an adaptation of the design used by [55] and [56]. Our design consists of a triangle shaped stainless-steel frame that supports a single high definition GoPro Hero 4 camera encased in an underwater housing (Fig 2). Each camera was provided with a backpack battery to extend its recording time to a minimum of 2 hours. Cameras were set to record at 60 frames per second/1080p resolution in wide field of view to maximize detection rates. All units had a baited arm to hold a perforated PVC bait container. A total of 1.5 kg of yellowfin tuna *(Thunnus albacares)* was used per each BRUVS during approximately 2 hr soak time (mean ± SD: 140 ± 13.4 min) since species accumulation in the open ocean after that time is less significant [33,55]. Once thawed, the bait was cut into approximately 5 cm pieces and lightly crushed once inserted into the perforated PVC container before each deployment. Drifting pelagic BRUVS were manually launched and retrieved from the vessel.

**Fig 2.**
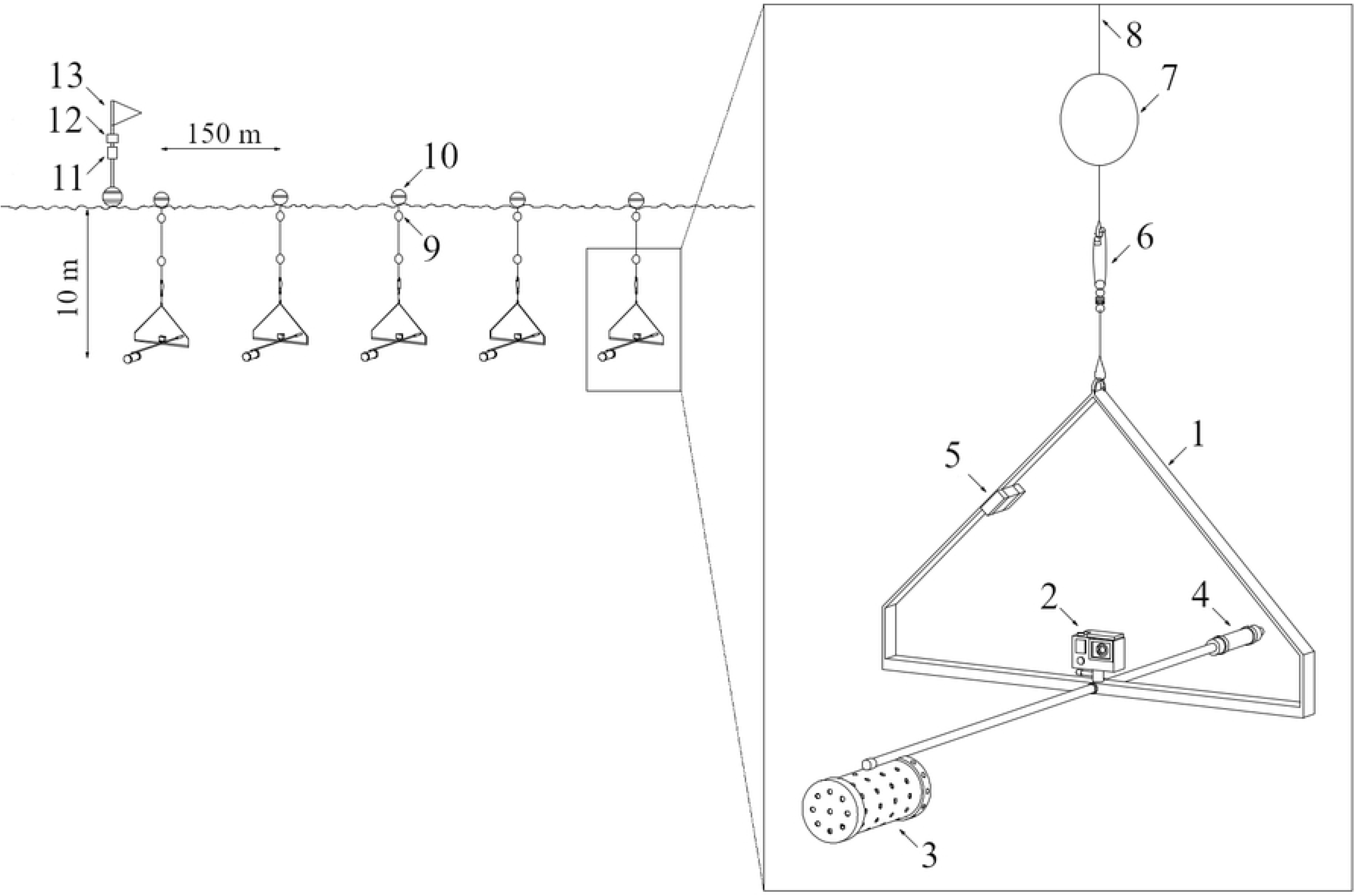
Diagram of a shallow (10 m depth) drifting-pelagic Baited Remote Underwater Video Stations (BRUVS) deployment. Numbers show closer details of a single BRUVS: 1) Triangle-shaped stainless steel frame; 2) GoPro camera; 3) Perforated PVC bait container; 4) Counterweight; 5) Temperature sensor; 6) Stainless steel tuna fishing swivel clip; 7) Small intermediate buoy; 8) 10 m or 25 m line; 9) Small surface buoy; 10) Big surface buoy; 11) Global Positioning System device (GPS); 12) Automatic Identification System (AIS) track device; 13) Flag

The drifting-pelagic BRUVS units were connected with a superficial tether line resembling a long-line system to maximize survey effort while minimizing the risk of loss and the amount of the tracking devices needed. Two long-lines with 5 pelagic-BRUVS separated 150 meters each were used during the expedition (Fig 2). We refer to each long-line as one deployment. To assess the effect of time of day on species richness and abundance, two deployments were simultaneously conducted during the morning (6 – 10 am) and two during the afternoon (1 – 3 pm) on the same seamount. Therefore, a total of four deployments (20 BRUVS) were conducted per site, except for North West Darwin and Las Gemelas 2, where only 2 deployments (10 BRUVS) were completed. Final soak times at each seamount ranged from 11.7 to 55.1 hours (mean ± SD: 32.5 ± 13.3 hours) (Table 1). To assess differences in species detectability between depths, simultaneous deployments were suspended at 10 m (shallow) and 25 m (deep) depth levels (Fig 2). All deployments entered the water upstream of each seamount to remove the effect of the vessel when the units drift over the sampling sites. The average distance left among simultaneous deployments was 1.6 ± 1.04 km (mean ± SD).

Differences in temperature between shallow and deep deployments per seamount were obtained attaching a temperature datalogger (ONSET Hobo Pendant^®^ UA) to each BRUVS unit. The drifting speed of each deployment was measured fitting each deployment with a Global Positioning System device (GPS). An Automatic Identification System track device (AIS) was also attached to each deployment to capture its position while drifting with the current (Fig 2). Missing temperature and drifting speed values for some BRUVS (8% and 6.7% respectively) were obtained calculating the average of those values between the two nearest stations. Temperature and drifting speed measurements of each deployment were averaged per seamount (Table 1). Concentrations of chlorophyll-A at each seamount were extracted from ERDDAPP open source data server using the *rxtracto* function from the *rerdappXtracto* package (Table 1). Additional information such as date, time, location and duration of each BRUVS was recorded on the field.

### Video and data analysis

The software EventMeasure (SeaGIS®) was used to analyze the video footage and calculate the relative abundance of each observed species. For relative abundance, we used the MaxN, which was defined as the maximum number of individuals from each species detected in a single frame of the video. The MaxN is a conservative estimate of the total number of individuals present in the deployment area because it avoids repeat counts of animals reentering the field of view and because only a proportion of the species present in the deployment area will positively respond to the bait plume by entering the camera’s field of view [57,58]. Species were identified to the lowest taxonomic level possible using local identification guides and expert knowledge if required. Species were classified in four ecological groups according to taxonomy and reported common sizes [59] into elasmobranchs, large teleosts (species with common total length > 1m), small teleosts (species with common total length < 1m) and other marine megafauna species (dolphins and turtles). Small teleost richness and abundance data were described, but not included in the statistical comparisons since their higher abundance could mask the distribution and abundance patterns of LPS such as elasmobranchs, large teleosts and other marine megafauna species which were pooled together during the analysis. LPS are of particular conservation concern because of targeted overfishing, high bycatch rates and declining population trends [60].

Differences in sampling effort were standardized across deployments and seamounts by dividing MaxN of each species obtained per BRUVS unit by soak time (effective time of BRUVS recording), expressed as MaxN·hr-^1^. BRUVS from the same long-line were separated 150 m from each other, and most likely were not independent. Therefore, we considered each deployment (one long-line with 5 connected BRUVS) as an independent replicate. The maximum MaxN·hr-^1^ per deployment was summed across seamounts and ecological groups for comparison purposes. Measures of dispersion of observed means were reported as mean ± standard deviation (SD).

Cumulative species richness curves were used to examine temporal accumulation of new species per group by BRUVS deployment and soak time. The order in which species were analyzed was randomized 999 times and the cumulative number of new species per BRUVS was counted for each randomization.

A non-parametric two tailed Wilcoxon Rank Sum test at 5% level of significance (α=0.05) was used to compare richness and abundance of LPS according to three different variables: time of deployment (morning vs afternoon), depth of deployment (shallow deployments vs deep deployments) and depth of seamount (shallow seamounts vs deep seamounts). Marine predators in the Azores and along the Indo-Pacific region showed a higher tendency to be associated with shallow seamounts (< 400 m) [38,61]. Therefore, we categorized seamounts shallower than 400m as shallow seamounts and seamounts deeper than 400m as deep seamounts (Table 1). The non-parametric Wilcoxon Rank Sum test was used because the normality assumption was not satisfied neither for LPS abundance (Shapiro-Wilk, p < 0.001) nor LPS richness (Shapiro-Wilk p = 0.001) and because comparisons where made between two independent samples [62]. Differences in the abundance of each ecological group according to these three variables were also tested using a Wilcoxon Rank Sum test and graphically represented. The Levene test was used to test homoscedasticity (equality of variances) between independent samples. Variances were not equal only when comparing LPS abundance between deep and shallow seamounts. Finally, a Wilcoxon Rank Sum test was also used to compare differences of environmental drivers (mean temperature, mean drifting speed and mean concentration of chlorophyll-a) between shallow and deep deployments, morning and afternoon deployments and shallow and deep seamounts in order to assess their potential effect on LPS distribution. All statistical tests were performed using R statistical packages (v 3.6.3). The function *wilcox.test* from the *MASS* package was used to obtain the statistics W value and the function *wilcox_test* from the package *coin* was used to calculate the Z value and the exact p value in all cases. To calculate the effect size (r) we divided the Z value by the square root of the sample size.

## Results

### Richness and abundance

We analyzed 347.5 hours of video footage from 32 deployments (150 BRUVS). Animals were detected on 97% of all deployments (N = 31) or 88% of individual BRUVS (N = 132), with the number of species per deployment ranging from 1 to 6 (mean ± SD: 2.7 ± 1.6 species). Overall, BRUVS detected 21 species, from which 14 species were teleosts (6 families), 3 sharks (3 families), 2 pelagic rays (2 families), 1 dolphin and 1 sea turtle (Table 2). Some individuals were only detected at the genus level *(Mobula* spp. and *Decaptreurs* spp.) and few small pelagic teleosts (56% of all sighted individuals) could not be identified due to their small size and/or low video resolution, hereafter referred as “unidentified”. Of all species detected 19% were threatened and 71% were non-threatened species, based on the International Union for the Conservation of Nature (IUCN) Red List assessments [63]. Only 5% of all identified species were classified as either data deficient or not evaluated (Table 2).

**Table 2.** Summary of occurrence, relative abundance (MaxN) and conservation status of all species recorded on baited remote underwater video stations (BRUVS) along Cocos Ridge seamounts.

| EG / Family | Species | Common name | N | % N | MaxN |  | MaxN·hr<sup>-1</sup><br>(mean ± SD) | Rank | Status |
| --- | --- | --- | --- | --- | --- | --- | --- | --- | --- |
|  |  |  |  |  | max | sum |  |  |  |
| Elasmobranchs |  |  |  |  |  |  |  |  |  |
| Sphyrnidae | *Sphyrna lewini* | Scalloped hammerhead | 11 | 34.4 | 60 | 162 | 6.3 ± 8 | H | CR |
| Carharhinidae | *Carcharhinus falciformis* | Silky shark | 5 | 15.6 | 2 | 7 | 0.6 ± 0.2 | M | VU |
| Alopiidae | *Alopias pelagicus* | Pelagic tresher shark | 3 | 9.4 | 1 | 3 | 0.4 | L | VU |
| Mobulidae | *Mobula spp.* | Mobula ray | 2 | 6.2 | 2 | 3 | 0.6 ± 0.3 | L | - |
| Dasyatidae | *Pteroplatytrygon violacea* | Pelagic stingray | 2 | 6.2 | 1 | 2 | 0.4 | L | LC |
| Total |  |  | 16 | 50 | - | 177 | 3.3 ± 6 | - | - |
| Large teleosts |  |  |  |  |  |  |  |  |  |
| Coryphaenidae | *Coryphaena hippurus* | Common dolphinfish | 14 | 43.8 | 28 | 134 | 4.6 ± 5.1 | H | LC |
| Istiophoridae | *Kajikia audax* | Sripped marlin | 5 | 15.6 | 1 | 5 | 0.5 ± 0.2 | M | NT |
|  | *Istiompax indica* | Black marlin | 1 | 3.1 | 1 | 1 | 0.4 | L | DD |
|  | *Istiophorus platypterus* | Sailfish | 1 | 3.1 | 1 | 1 | 0.4 | L | LC |
| Scombridae | *Thunnus albacares* | Yellowfin tuna | 2 | 6.2 | 3 | 4 | 0.8 ± 0.5 | L | NT |
| Alepisauridae | *Alepisaurus ferox* | Long snouted lancetfish | 1 | 3.1 | 1 | 1 | 0.4 | L | LC |
| Total |  |  | 19 | 59 | - | 146 | 2.9 ± 4 | - | - |

| Small teleosts |  |  |  |  |  |  |  |  |  |
| --- | --- | --- | --- | --- | --- | --- | --- | --- | --- |
| Carangidae | *Naucrates ductor* | Pilot fish | 16 | 50 | 37 | 105 | 2.7 ± 3.4 | H | LC |
|  | *Caranx caballus* | Green Jack | 4 | 12.5 | 500 | 553 | 55.4 ± 95.2 | M | LC |
|  | *Seriola peruana* | Fortune jack | 3 | 9.4 | 12 | 19 | 2.6 ± 2.2 | M | LC |
|  | *Seriola rivoliana* | Pacific amberjack | 1 | 3.1 | 6 | 6 | 2.5 | L | LC |
|  | *Decapterus spp.* | Scads | 1 | 3.1 | 1 | 1 | 0.4 | L | NE |
| Monacanthidae | *Aluterus monoceros* | Unicorn filefish | 2 | 6.2 | 3 | 4 | 0.7 ± 0.5 | L | LC |
| Coryphaenidae | *Coryphaena equiselis* | Pompano dolphinfish | 1 | 3.1 | 4 | 4 | 1.8 | L | LC |
| Scombridae | *Sarda orientalis* | Striped bonito | 1 | 3.1 | 3 | 3 | 1.2 | L | LC |
| Unidentified | Unidentified | Unidentified | 31 | 96.9 | 1000 | 1301 | 17.9 ± 74.9 | H | - |
| **Total** |  |  | **30** | **94** | **-** | **1996** | **13.9 ± 59** |  |  |
| Other marine megafauna |  |  |  |  |  |  |  |  |  |
| Delphinidae | *Tursiops truncatus* | Bottlenose dolphin | 9 | 28.1 | 10 | 33 | 1.8 ± 1.7 | M | LC |
| Cheloniidae | *Chelonia mydas agassizi* | Black pacific sea turtle | 1 | 3.1 | 1 | 1 | 0.4 | L | EN |
| **Total** |  |  | **9** | **28** | **-** | **34** | **1.7 ± 2** |  |  |

Species are organized by families and ecological groups (EG). N: number of BRUVS deployments with a species. Relative abundance is expressed as (i) MaxN: maximum number of individuals of a species per deployment (one deployment with 5 connected BRUVS is considered an independent sample) and (ii) MaxN·hr<sup>-1</sup>: MaxN divided by soak time (hours). Species are ranked based on the frequency of occurrence: H – high (>30% of deployments); M – medium (10-30% of deployments); and L – low (<10% of deployments). Conservation status of each species (DD – data deficient, LC – least concern, NT – near threatened, VU – vulnerable, EN –
endangered, and CR – critically endangered) is presented based on current IUCN Red List [63].

Small teleosts were the most abundant (13.50 ± 58.2 ind hr^−1^) and commonly sighted group (94% of all deployments). Large teleosts and elasmobranchs occurred at 59% and 50% of all deployments, respectively, and showed similar relative abundances (large teleosts: 2.9 ± 4.4 ind hr^−1^; elasmobranchs: 3.3 ± 6.1 ind hr^−1^). Other marine megafauna such as dolphins and sea turtles were the least sighted and abundant group, occurring at 28% of all deployments (1.7 ± 1.7 ind hr^−1^) (Table 2).

The scalloped hammerhead shark *(Sphyrna lewini)* clearly dominated the elasmobranch group, representing 91.5% of the group’s MaxN. Furthermore, this species showed the highest frequency of observation among all elasmobranchs (34.4% of all deployments) (Table 2). The common dolphinfish or mahi-mahi *(Coryphaena hippurus)* was the most abundant large teleost (91.8% of the group’s MaxN) and the most frequently observed among all large teleosts (43.8% of all deployments) (Table 2). Among the small teleosts, the green jack *(Caranx caballus)* was the most abundant species (79.6% of the group’s MaxN). However, the pilot fish (*Naucrates ductor*) was the most frequently observed species (50% of all deployments) (Table 2). The bottlenose dolphin *(Tursiops truncatus)* was the most abundant among other marine megafauna species (97% of the group’s MaxN) and the most frequently observed (28% of all deployments) (Table 2). Unidentified species were not considered in the ranking since their associated values were summed across several unidentified species.

The slope of the species accumulation curve indicated that the amount of species for all groups recorded increased gradually with soak time and number of BRUVS deployments; however, some differences were detected between groups (Fig 3). Although still increasing, the species accumulation curve for small teleosts nearly reached an asymptote after 180 min of soak time, whereas elasmobranchs and other marine megafauna species showed signs of stabilization after 90 min of soak time. The curve for large teleosts continued to increase at a maximum soak time of 140 min (Fig 3A). Species richness accumulation curves increased at a faster rate for small teleosts and elasmobranchs (there was a sharp increase during the first 20 minutes), followed by large teleosts and ultimately by other megafauna species at a much slower rate (Fig 3A). Finally, all groups showed an increasing trend of species accumulation over the number of BRUVS deployments (Fig 3B).

**Fig 3.**
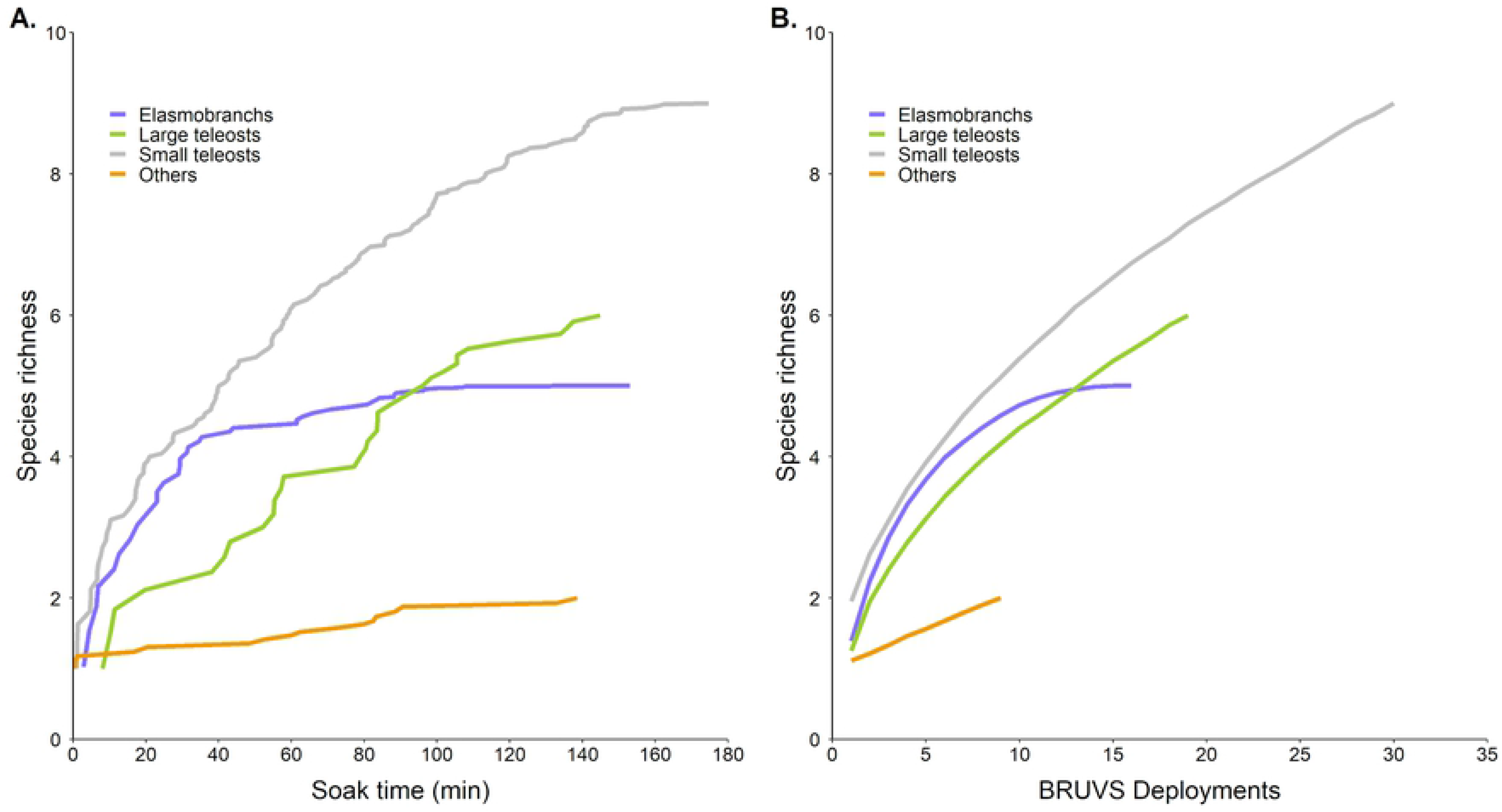
Species accumulation curves per ecological group. (A) Species richness of each ecological group over cumulative soak time in minutes. Soak time is defined as the effective recording time of Baited Remote Underwater Video Stations (BRUVS). (B) Species richness of each ecological group over the number of BRUVS deployments. Each deployment includes 5 BRUVS and is treated as an independent sample.

### Spatial distribution across seamounts

The highest richness and relative abundance of LPS were found at West Cocos (richness: 3.8 ± 1.5 species; MaxN hr^−1^: 12.8 ± 6.2 ind hr^−1^) and Paramount (richness: 2.5 ± 1 species; MaxN hr^−1^: 9.1 ± 9.9 ind hr^−1^) seamounts, whereas the lowest values were found at Medina 3 (richness: 0.5 ± 0.6 species; MaxN hr^−1^: 0.2 ± 0.3 ind hr^−1^) and NW Darwin (richness: 0.5 ± 0.7 species; MaxN hr^−1^: 0.2 ± 0.4 ind hr^−1^) seamounts (S1 Table).

The number of LPS ranged from 1 species at NW Darwin to 8 species at West Cocos (Fig 4A; S1 Table). Elasmobranchs were detected at all seamounts except for Medina 1 and NW Darwin, whereas large teleosts were not detected at Paramount and Las Gemelas 2. Other marine megafauna species were only reported at seamounts close to Cocos Island including Las Gemelas seamounts, West Cocos and East Cocos (Fig 4A). The highest elasmobranch richness was reported in Paramount, whereas the highest richness of large teleosts and other marine megafauna species were reported at West Cocos (Fig 4A).

**Fig 4.**
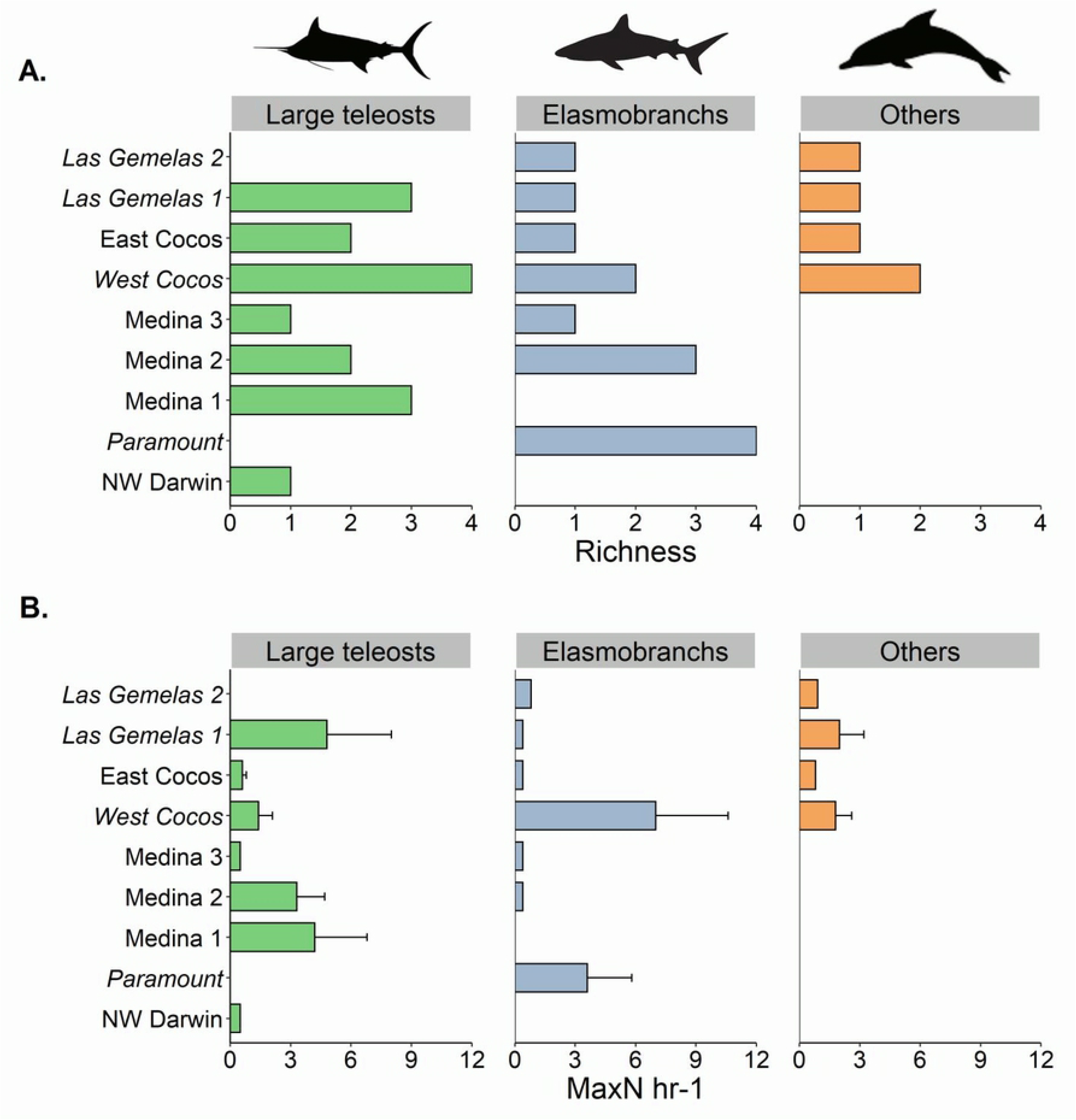
Comparison of (A) species richness and (B) relative abundance (MaxN hr^−1^) among ecological groups. Seamounts are ordered from bottom to top according to the distance from Galapagos Islands (bottom) to Cocos Island (top). Error bars represent the 95% confidence intervals. Shallow seamounts (<400 m) are shown in italics.

Elasmobranchs were more abundant at West Cocos and Paramount seamounts (Fig 4B) where large schools of *S. lewini* of up to 40 and 60 individuals were observed in a single deployment, respectively (S1 Table). Few individuals of *S. lewini* were also detected in Medina 3 and Gemelas 2 (S1 Table). Large teleosts were more abundant at Las Gemelas 1 and Medina 1 (Fig 4B), where our BRUVS reported schools of 27 and 28 individuals of *C. hippurus* in a single deployment, respectively (S1 Table). However, the highest abundance of tunas (*Tunnus albacares*) and billfishes (*Istiompax indica, Istiophorus platypterus* and *Kajikia audax*) was reported at West Cocos (S1 Table). Other marine megafauna species were more abundant at West Cocos and Las Gemelas 1 (Fig 4B) where our BRUVS recorded groups of up to 6 and 10 *T. truncatus*, respectively (S1 Table). The black sea turtle *Chelonia mydas agassizi* and *T. truncatus* were observed only at seamounts close to Cocos Island (S1 Table).

The Wilcoxon Rank Sum Test showed a significant effect of seamount depth on LPS relative abundance (W= 49.5, Z = 2.3, p < 0.05, r = 0.4) and LPS richness (W = 43, Z = - 3.3, p < 0.001, r = 0.6). Relative abundance of LPS was significantly higher at shallow seamounts (mean ± SD = 8.2 ± 7.9 ind hr^−1^) compared to deeper ones (mean ± SD = 2.6 ± 4.3 ind hr^−1^). Mean richness of LPS at shallow seamounts (mean ± SD = 2.6 ± 1.3 species) was also higher compared to deep seamounts (mean ± SD = 1.2 ± 0.8 species). When evaluating differences in environmental variables between types of seamounts, we found that only chlorophyll-a (W = 214, Z = 3.4, p < 0.001, r = 0.6) and drifting speed (W = 231, Z = 4, p < 0.001, r = 0.70) were significantly lower in shallow seamounts.

Differences between morning and afternoon deployments were not significant either for LPS abundance (W = 131.5, Z = 0.1, p = 0.9, r = 0.02) or LPS richness (W = 139, Z = 0.4, p = 0.7, r = 0.07). Furthermore, none of the environmental variables were significantly different between morning and afternoon deployments. Although temperature was significantly higher at shallow deployments (mean ± SD = 28.8 ± 0.8 °C) compared to deep deployments (mean ± SD = 26.9 ± 1.7 °C), we didn’t find significant differences on LPS abundance (W = 138.5, Z = 0.4, p = 0.7, r = 0.07) nor LPS richness (W = 142.5, Z = 0.6, p = 0.6, r = 0.1) between camera depth levels. None of the other environmental drivers were significantly different between shallow and deep deployments.

When comparing relative abundance among ecological groups, significant differences were only found between deep and shallow seamounts for elasmobranchs (W = 38.5, Z = −3.55, p < 0.001, r = 0.63) and other megafauna species (W = 59, Z = −3.21, p < 0.001, r = 0.58) (Fig 5).

**Fig 5.**
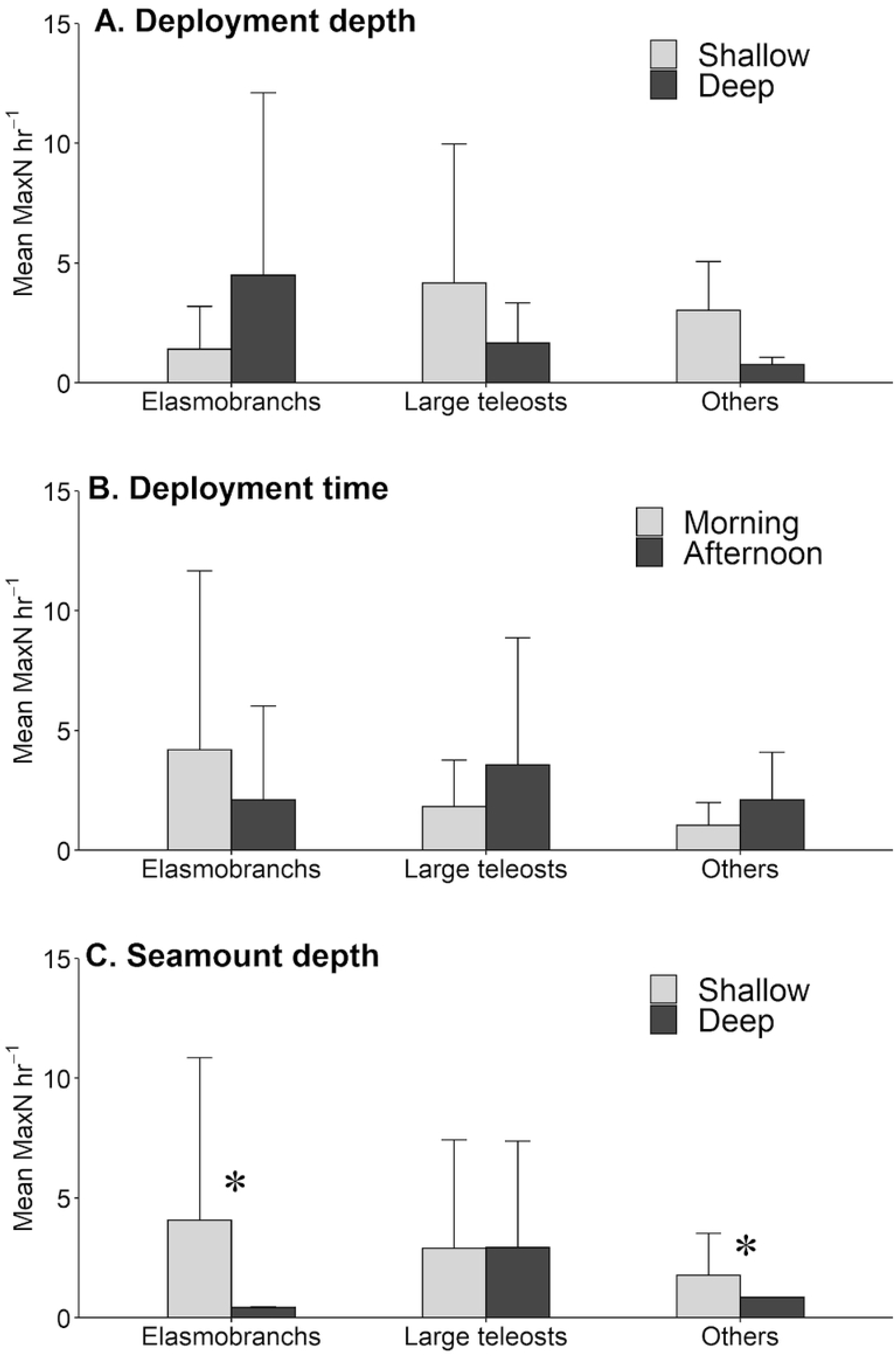
Comparison of mean relative abundance (MaxN hr^−1^) among ecological groups according to three different variables. : A) shallow (10m) and deep (25m) deployments; B) morning (6-10am) and afternoon (1-3pm) deployments; C) shallow (<400m) and deep (>400m) seamounts. Significant differences among levels are shown with (*).

## Discussion

### The usefulness of drifting pelagic-BRUVS

Understanding how pelagic species assemblages are influenced or shaped by seamounts is of critical importance to successfully manage and protect open-ocean ecosystems [64]. Our study showed that drifting pelagic BRUVS are an effective, accessible and non-extractive technique to monitor LPS in oceanic environments, highlighting its potential to guide management and conservation actions in the ETP.

Despite the reduced spatial and temporal replication of this study, our drifting-pelagic BRUVS showed a high frequency of species detection (97%) compared to other studies using this same technique in topographically complex pelagic environments of Australia [37] and the Indo-Pacific region [38], where species detection was 82% and 87%, respectively. However, the number of independent replicates in this study was lower (n=31 deployments) compared to [37] and [38] were 51 and 1041 pelagic BRUVS were deployed, respectively. Futhermore, treating five connected individual BRUVS as an independent replicate in this study may have also increased the species detectability in comparison with these other studies where each BRUVS unit was considered an independent replicate. In any case, if we consider each BRUVS as an independent replicate, species detectability remains high (88%). Despite more replicates and similar sampling protocols (or experimental/field study designs) are needed to establish meaningful comparisons among regions, our results demonstrate that threatened and vulnerable species with high commercial value (in many cases) were repeatedly detected by our drifting-pelagic BRUVS at shallow depths along unprotected waters of the marine corridor between Cocos and Galapagos Islands.

All shark species detected by our drifting pelagic BRUVS are listed by the IUCN Red List as either Critically Endangered (*S. lewini*) or Vulnerable (*Carcharhinus falciformis* and *Alopias pelagicus*) [63]. These species represent one of the main components of the elasmobranch bycatch associated with long-line and gill-net fisheries in the ETP [25,26,65,66], mainly because of the high commercial value of their fins for the shark fin trade worldwide [67,68]. As a result, some studies have reported large population declines of *S. lewini,* silky sharks (*C. falciformis*) and thresher sharks (*A. pelagicus*) in the ETP [19,21,69]. For example, the abundance of *S. lewini* appears to have declined by 50% in Galapagos Islands since 1980 [70], by 45% in Malpelo Island since 2000 [21] and by 50% in Cocos Island since 1990 [19] suggesting that protection from oceanic MPAs in the ETP is likely not enough to prevent shark declines, particularly of wide-ranging migratory species.

It is important to highlight that our BRUVS captured groups of up to 60 and 40 individuals of *S. lewini* over two different seamounts providing the first visual evidence of the schooling behavior of this species even in open-water ecosystems. This tendency of *S. lewini* to form large aggregations, presumably near shallow seamounts, makes it highly vulnerable to pelagic-longline and purse-seine fisheries [25,71,72]. The schooling behavior of *C. hippurus* and *T. truncatus* may also have contributed to their higher abundance among other large teleosts and marine megafauna species, respectively. Although teleost fishes are R-selected species (high growth rates, early maturity, high fecundity and many offspring), and therefore, more resilient to overfishing than elasmobranchs [73], their heavy commercial exploitation by several coastal nations of Latin America and the poor fisheries management frameworks make these species a conservation concern [25,26]. For example, *C. hippurus* is commonly captured by artisanal pelagic longline fisheries in the ETP, and represents 65% of the estimated landings in Ecuador [26,74] and Costa Rica [65]. Marine turtles and cetaceans are also captured as bycatch species in longline fisheries operating within the ETP [25,75,76]. Overall, our results demonstrate the utility of using drifting-pelagic BRUVS to survey highly mobile species in pelagic environments.

Despite baited camera surveys have been positively validated against extractive methods such as trawling and longlines [39,77], BRUVS also have limitations [36,78]. For example, bait plume dispersion is complex and dynamic which makes surveyed area unknown [58]. Also, bait responses behaviors may lead to sampling biases towards larger mobile species, yet in this scenario, they were the target group of our study. Since our BRUVS were deployed in offshore clear water (> 20 meters), visibility didn’t compromise our data. Although our cameras recorded under a wide field of view (220 degrees), there is still a portion of the surrounding area that was not sampled effectively, and therefore counts of relative abundance could be underestimated. This problem can be solved by using 360° cameras that allow a full field of view around each BRUVS, but it will also increase surveying costs and time of analysis. Despite the above mentioned limitations, drifting-pelagic BRUVS generate relevant ecological data of apex predator guilds that are typically cryptic, increasingly exposed to anthropogenic mortality and of high conservation and commercial value without posing a threat to targeted species [36].

### Towards a regional standardization of drifting pelagic-BRUVS

Some efforts have been made to develop operation procedures that ensures data standardization in studies using benthic [79] and pelagic BRUVS [36,78]. However, most of the studies available were conducted in Australian waters, and did not account specific environmental and oceanographic conditions inherent of the ETP such as the interannual variability of the physical environment due to El Niño Southern Oscillation (ENSO), low salinity surface waters, moderate productivity, large hypoxic waters and shallow thermoclines [49,80]. A standardized protocol for the ETP would allow spatio-temporal comparisons of abundance and richness patterns among different locations in the region. Our results can be used as a first approach to guide future studies using drifting-pelagic BRUVS in the region.

Species accumulation curves in this study indicate that with higher survey effort more species from all groups would be detected. Survey effort in open-water ecosystems is expected to be higher than in demersal habitats due to lower densities of organisms [33]. However, our results reveal a variation at a group level in species accumulation rates that is important to consider. For example, at least 90 min were needed to record a representative sample of apex predators such as elasmobranchs and cetaceans, whereas soak times up to 180 minutes were necessary to increase species detection of small and large teleosts. The early presence of predators around BRUVS and the higher attraction of predators to the bait could have affected the presence of smaller species (but see 79). These results coincide with those reported by [78] where the slope of the curve for pelagic species in Western Australia was reduced after 90 minutes of soak time with a trend still increasing at a reduced rate at 180 minutes. Overall, an average of 120 minutes soak times was identified by [78] as the optimal soak time to sample pelagic environments. However, midwater BRUVS in [78] were deployed while anchored on the seafloor at 35m depth between 1 and 2 km from the reef slope of the north-west cape of Western Australia. In this study, instead, we used drifting-pelagic BRUVS to survey fully pelagic habitats over sea bottom depths of at least 1000 m at minimum and maximum distances of 27.5 km and 327 km from the nearest oceanic islands, respectively. Therefore, the optimal soak time in pelagic environments is likely to be higher as the distance from continental shelves and sea bottom depth increase.

The rapid increase of species accumulation in the first 15-20 min shown by elasmobranchs and small teleosts in this study has been also reported by [78] and [37], although differences at the group level were not evaluated in any of these studies. Differences in species accumulation rates among groups may be related to differences in species richness by group, the specific responses of each group to the different BRUVS stimuli (olfactory, visual or auditory) and to the schooling behavior of each group [40]. Therefore, we recommend considering such differences when selecting optimal survey effort using drifting-pelagic BRUVS.

The movement capacity of large pelagic species [58], and the uncertainty associated with bait plume dispersal and sampling area during BRUVS studies [36], has caused debate about the minimum distance necessary to guarantee independence between replicates [36,78,79]. Accurate current speed measurements and bait dispersion models, although complex, are necessary to determine such minimum distance [58]. For example, a maximum bait plume extension of 920 m was calculated at maximum current speed of 0.34 m s^−1^ and 45 min of soak time [58]. In the absence of such data, we recommend leaving at least 1 km as a conservative distance between independent drifting-pelagic BRUVS deployments to reduce potential biases associated with pseudo-replication.

In this study, neither the depth level of the cameras nor the time of deployment had a significant influence on the abundance and richness of LPS captured by our BRUVS. In the ETP, the low levels of dissolved oxygen concentrations as a result of high productivity near to the surface create hypoxic conditions and a shallow thermocline that limit habitat suitability of LPS at extremely shallow depths up to 25 m [49,80]. Therefore, it is not surprising that no significant differences in LPS richness and relative abundance were found between shallow and deep BRUVS deployments since variations of environmental parameters are expected to occur below the thermocline. However, our data showed different patterns between groups that might be important to consider for future studies. For example, elasmobranchs were more abundant at deep deployments (25m) during the morning whereas large teleosts were more abundant at shallow deployments (10m) during the afternoon. Such variations can be attributed to species-specific behavioral adaptations such as foraging, predator avoidance or reproductive aggregations [82]. Furthermore, seamount production can be supported by a variety of energy inputs, and therefore, different responses at each species or trophic group level can be expected [83]. The fact that elasmobranchs were more abundant at deeper deployments could be associated with the thermocline at the same depth level [49,80]. This pattern has been previously reported for sharks surveyed at Malpelo Island, an oceanic island of the ETP [17]. Although in this study we did not detect the depth of the thermocline, our data showed significantly lower temperatures in deep deployments compared to shallow ones, which could be affecting the distribution of LPS.

Since environmental drivers in the ETP respond to seasonal variations [18,50], we recommend exploring the effect of depth level and time of deployment on LPS at larger spatial and temporal scales (at least twice a year covering dry and rainy season). Additional variables such as dissolved oxygen [80], ocean currents [58,84], prey availability [61] and fishing effort [38] might also pose a significant influence on LPS and therefore should be considered as potential drivers. Although we did not explore all these parameters in depth, our results can serve as a baseline to incorporate environmental drivers and their effect on LPS within the ETP.

### Large pelagic species at seamounts along the Cocos Ridge

Our drifting-pelagic BRUVS detected LPS in the vicinity of all surveyed seamounts in an open ocean environment where species richness is naturally low [85]. The evidence of the seamount effect on aggregating pelagic fauna has been demonstrated in a numerous studies and for a wide variety of species including zooplankton [84,86], teleosts [61,87–90], sharks [38,91,92], dolphins [93] and sea birds [94]. Furthermore, seamount-related fisheries represent a significant proportion of the total high seas fish catch highlighting the importance of these unique topographic features as hotspots for pelagic biodiversity [6,60,95]. For example, bathymetry and seamount presence were identified as one of the most important predictors of illegal fishing activities inside Cocos Island [96]. Despite all this evidence, more research effort along the Cocos Ridge is needed to elucidate if pelagic assemblages in the area are positively associated to seamounts presence.

Our results showed significantly higher richness and abundance of LPS at shallow seamounts (< 400m) compared to deeper ones (> 400m) suggesting that seamount depth could be an important parameter to identify aggregations of LPS along the Cocos Ridge. The influence of seamount depth on predator species have been previously reported at the Azores Islands [6] and the Indo-Pacific region [38] where remote seamounts shallower than 400m and 500m respectively showed a significant aggregation effect on predator species. Higher fishing catches have been also reported to occur at seamounts shallower than 400m compared to deeper ones [84]. The most common explanation given to this pattern is that shallower abrupt topographies block the vertically migrating zooplankton and micronekton that ascend at night and accumulates between the photic zone and 400m providing sufficient food to maintain higher trophic levels and commercial fisheries [84,97].

The influence of seamount depth on LPS could explain the highest abundance and richness of LPS found at West Cocos (283 m) and Paramount (188 m) and the lowest abundance and richness of LPS found at NW Darwin, the deepest surveyed seamount (~1200 m). Furthermore, large aggregations of *S.lewini* were only found at West Cocos and Paramount. Therefore, we recommend a special emphasis on both seamounts for future studies in the region. Measuring fishing pressure at these and other seamounts is of paramount importance in order to elucidate their degree of disturbance and consequently their conservation priority [24].

Since Las Gemelas seamounts are among the shallowest surveyed seamounts, we would have expected higher concentrations of sharks and higher richness of LPS. However, the reported abundance of LPS in Las Gemelas seamounts was mainly attributed to *C. hippurus* and in general, LPS abundance was relatively low compared to the high abundance of smaller teleosts (S1 Table). These results coincide with [98] which found a higher fish biomass at Las Gemelas compared to Cocos Island but a much lower abundance of large predatory fishes. One possible explanation is that seamounts closer to MPAs receive greater fishing pressure since they are more accessible to humans and they are of special interest to fishers because they benefit from spillover effects [99]. Global Fishing Watch data showed how, in certain seasons, fishing fleets in the ETP concentrate their operations surrounding the protected waters of Cocos Island [100] potentially affecting Las Gemelas seamounts located at only 50 km from the MPA. Although Las Gemelas seamounts are managed as a no-take zone within the Seamounts Marine Management Area since 2011, enforcement capacity in this area is very limited and therefore illegal fishing still remains a major threat [100,101]. Given the diverse and rich assemblages of invertebrates carpeting the bottom habitats and the high biomass of prey fishes in Las Gemelas seamounts [98,102], is very likely that large predators could rapidly increase if fishing activities were effectively prohibited.

Our data showed that elasmobranchs and other marine megafauna species (mainly dolphins) are the main groups affected by seamount depth, while no significant difference of large teleosts abundance was found between shallow and deep seamounts. This could explain the low abundance of elasmobranchs at deep seamounts such as Medina 1, Medina 2, Medina 3, East Cocos and NW Darwin. However, there is also the possibility that our BRUVS located at a shallow depth from the surface failed at capturing pelagic life at deeper levels.

Chlorophyll-a is known to increase the availability of food and therefore pelagic species diversity and abundance of large predators [103–105]. However, in this study we found significantly lower concentrations of chlorophyll-a at shallow seamounts where abundance of LPS was significantly higher. Similarly, strong currents are usually associated with enhanced primary production and therefore higher abundance predator species [84]. However, our results showed significantly lower drifting speeds at shallower seamounts where abundance of LPS was significantly higher. This may imply that there are other environmental factors not measured in this study that have a greater influence on LPS distribution along the Cocos Ridge. However, is more likely that a higher survey effort including sampling along the water column and at different distances from seamounts is necessary to understand the environmental variations at each seamount and its effect on LPS.

## Conclusion

Our study represents the first attempt at characterizing the spatial distribution and relative abundance of LPS along underwater seamounts of the Cocos Ridge, providing a first insight on how pelagic communities are structured outside the protection limits of two of the most important oceanic MPAs of the ETP. Shallow seamounts presented significantly higher abundances of LPS, and thus may represent ecologically important refugees for LPS in the ETP. However, survey effort in this study was not enough to demonstrate a positive association between seamounts and LPS. Future research is necessary to assess the potential of seamounts as high-priority conservation areas to prevent threatened species from further declines. Our results show a promising potential of the drifting-pelagic BRUVS to survey LPS in pelagic ecosystems of the ETP. This study might serve as an important reference for future studies in the region using this technique.

## Acknowledgments

This study would not have been possible without the support and collaboration of the Área de Conservación Marina Cocos, the Galapagos Science Center and Dirección del Parque Nacional Galapagos. We would like to thank I. Chaves, T. Araya and J. Valerio who supported us with the video analysis. We also would like to thank all the crew from the Plan B vessel, the research boat used during the expedition, for their constant support during the field work. We are also grateful with Ignasi Agustí Fuster for his collaboration on the Baited Remote Underwater Video Station diagram presented on this paper. This study was conducted under research permit PC-24-18 issued by the Galapagos National Park Directorate”

## Supporting information

**S1 Table. Relative abundance (MaxN hr^−1^) and richness of large pelagic species (LPS) by seamount.** Relative abundance is expressed as MaxN hr^−1^ (maximum number of individuals of a species recorded on a single deployment standardized by soak time). MaxN hr^−1^ Species are abreviated as: AF) *Alepisaurus ferox,* AP) *Alopias pelagicus,* CF) *Carcharhinus falciformis*, CM) *Chelonia mydas agassizi*, CH) *Coryphaena hippurus*, II) *Istiompax indica*, IP) *Istiophorus platypterus*, KA) *Kajikia audax*, M) *Mobula* spp., PV) *Pteroplatytrygon violacea,* SL) *Sphyrna lewini,* TA) *Thunnus albacares* and TT) *Trusiops truncatus*. The MaxN hr^−1^ at a group level for the small teleost species (ST) is also presented. The sum and the mean of LPS MaxN hr^−1^ and species richness are also presented per seamount. Seamounts are ordered according to the sum of MaxN hr^−1^. Shallow seamounts (<400 m) are shown in italics.

